# MagC, magnetic collection of ultrathin sections for volumetric correlative light and electron microscopy

**DOI:** 10.1101/526137

**Authors:** Templier T.

## Abstract

The non-destructive collection of ultrathin sections onto silicon wafers for post-embedding staining and volumetric correlative light and electron microscopy traditionally requires exquisite manual skills and is tedious and unreliable. In MagC introduced here, sample blocks are augmented with a magnetic resin enabling remote actuation and collection of hundreds of sections on wafer. MagC allowed the correlative visualization of neuroanatomical tracers within their ultrastructural volumetric electron microscopy context.

## Introduction

The ultrathin physical ablation of sample blocks is a prerequisite for volumetric biological electron microscopy (EM). The destructive methods serial block face^1^ and focused ion beam EM^2^ enable serial access to the sample in its whole depth only very briefly and inside the vacuum chamber of a specialized scanning EM, prohibiting (re-)imaging of permanently destructed portions, liquid treatments such as heavy-metal poststaining or immunostaining^3^, fluorescent light microscopy (LM)^4^, and various nanoscale imaging techniques^5^.

The automated non-destructive tape-based ablation method ATUM^6^, that greatly benefited volumetric EM^7^, provides sections onto silicon wafers but with a low packing density (about 200 per 100 mm diameter wafer), through an intermediate tape, and after manual gluing onto a wafer. Carbon-coated Kapton tape suffers from strong autofluorescence preventing fluorescence microscopy, from scratches impairing EM imaging^8^, and from the difficulty to be uniformly carbon-coated^8^ (a necessary step to avoid charging during imaging). Recently introduced carbon-nanotube tapes^8^ solve most of these issues, though they require a custom device for reel-to-reel plasma hydrophilization and manual grounding of all cut tape stripes with conductive tape on top.

The invention of MagC was motivated by the wish instead to collect sections directly onto silicon wafers for excellent fluorescent LM and EM imaging conditions without tape-related issues and at high packing density for convenient bulk staining procedures with liquids and uninterrupted imaging in automated LMs and EMs, including next-generation multibeam EM^9^. In MagC, a piece of resin containing superparamagnetic nanoparticles is glued onto a polymerized sample block so that all cut sections carry magnetic material. Remote magnetic actuation then allows the agglomeration of floating sections in the center of a large bath attached to a diamond knife until they are deposited onto an underlying silicon wafer. Finally, the order of the sections is retrieved computationally after section collection. Two volumetric correlative LM-EM data sets of connectomics-grade brain tissue are presented here.

## Results and discussion

### 1. Magnetic resin

Magnetic epoxy-based resin containing 8% (w/w) iron oxide superparamagnetic nanoparticles^10^ was produced for remote actuation. The resin also contained fluorescent polymer beads for post-collection section order retrieval, and a fluorescent dye to ease section segmentation (Fig. 1a, S2, S4). A piece of this resin was glued with an epoxy usually used for EM studies (durcupan) to a sample block of interest, with the help of a small mechanical device (Fig. S1) to maintain the blocks in position during curing in an oven. The resulting blocks were trimmed, and a second piece of resin (the “dummy”), consisting of a piece of heavy-metal stained and resin-embedded brain tissue, was glued to the magnetic resin to enhance cutting quality. The final block assembly was trimmed for ultrathin sectioning (Fig. 1a).

**Figure 1.**
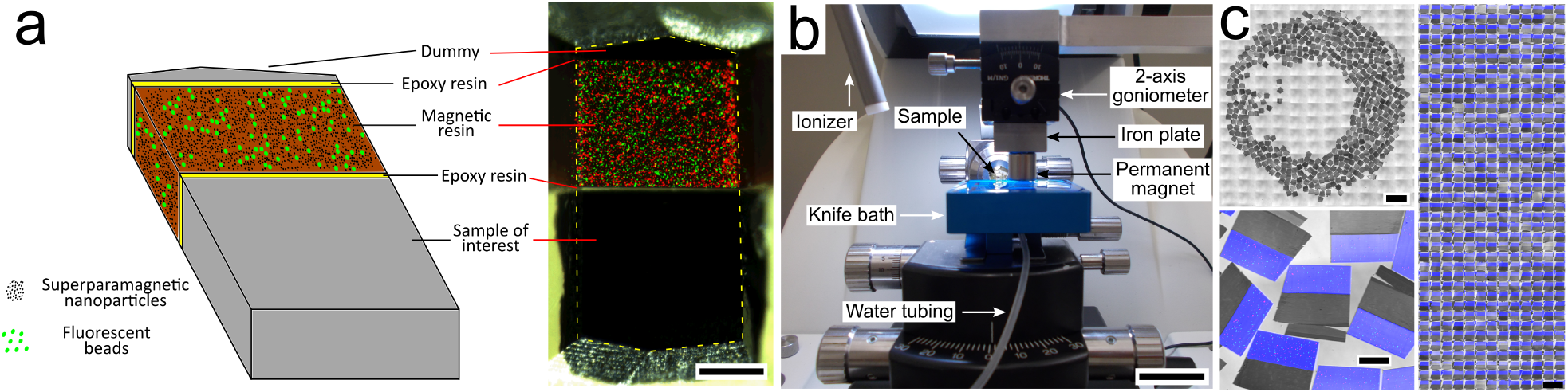
Magnetic augmentation and collection of sections on silicon wafer. **a**, Augmentation of a polymerized sample block with resin containing superparamagnetic nanoparticles (for remote magnetic actuation) and fluorescent beads (for section order retrieval). **b**, Setup for MagC: a diamond knife with a large bath and a mobile overhanging magnet. **c**, 507 consecutive ultrathin sections collected on a silicon wafer: wafer overview, close-up (merge of whitefield and 3 fluorescent channels: blue-Coumarin, green,red-fluorescent beads) and montage of all sections. Scale bars: **a**-200 μm, **b**-2 cm, **c**-2mm, 200 μm, 1 mm

### 2. Sectioning

A custom diamond knife was built with an enlarged bath to let many hundreds of sections float at the water surface (Fig. 1b). A hole was drilled in the bottom to fill and empty the bath with a motorized syringe pump. A piece of silicon wafer was immersed in the bath and was slightly tilted compared to the water level (about 2 degrees) to avoid accumulation of water surface dust in the center of the wafer at the end of the water withdrawal. After alignment of the knife and cutting a few sections, automatic sectioning was started and was left uninterrupted until the last section cut. A ionizer whose tip was placed close to the diamond knife created a very soft air current that gently detached sections from each other every few sections without impairing the cutting process. The sections floated freely at the water surface.

### 3. Magnetic collection

To collect the floating sections after the sectioning, a permanent magnet (cylindric, 15 mm diameter x 8 mm) was placed above the water surface with a 1 mm air gap (Fig. 1b). A few sections (about a dozen) that were slightly sticking to the walls of the bath were gently detached with an eyelash. The magnet, actuated by a robotic arm, scanned the water bath surface describing a snake path. At the end of the scan, the sections were accumulated in the center of the bath. Water was then withdrawn with a motorized syringe pump, while maintaining the 1 mm air gap by lowering the magnet with manual robotic control. Two small heating pads placed below the bath were turned on when the water level reached the level of the substrate.

The elevated wafer temperature generated by the heating pads (about 40 degrees) accelerated the evaporation of the water left at the wafer surface and avoided the formation of wrinkles in the deposited sections. The wafer was finally placed on a hot plate at 50 degrees for 30 minutes. I report here on two collected wafers of 507 (data set 1, Fig. 1c) and 203 consecutive 50 nm thick sections (data set 2, Fig. S2).

### 4. Order retrieval

The serial order was lost during the sectioning and had to be retrieved. After low-resolution (5x) reflection whitefield and fluorescent imaging of the wafers, the location and orientation of the sections was semi-automatically inferred. After calibration of 4 landmarks, medium resolution (20x) fluorescent imaging was automatically performed on the magnetic portion of each section. The cloud of fluorescent beads (2 μm mean diameter) contained in the magnetic resin was revealed in this imagery. Since the section thickness (50 nm) was smaller than the diameter of the beads, each bead was visible in at least a dozen of consecutive sections, so that pairwise similarity of all sections could be computed. Solving a traveling salesman problem on the graph of pairwise similarities retrieved the serial order (as confirmed later manually with EM) with only a single error when using a high concentration of beads (1% w/w, Fig. 2a). A lower concentration yielded more errors (0.2% w/w, Fig. S3). With the same methodology, the serial order could also be retrieved using the brain tissue EM imagery (Fig. 2a). Note that the order retrieval with fluorescent beads contained in the magnetic resin does not depend on the processed sample, which makes MagC suitable for collecting samples that for example would not show sufficient information for order retrieval by LM or EM.

**Figure 2.**
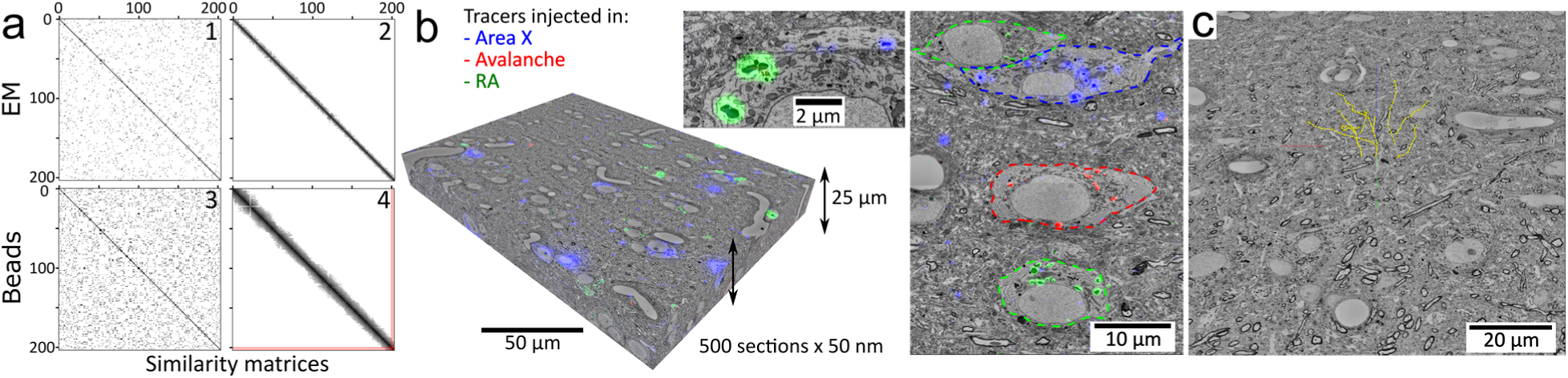
Volumetric correlative LM-EM with MagC collected sections. **a**, Section order retrieval on data set 2 (1% fluorescent beads) obtained with EM imagery (panels 1,2 show the pairwise similarity matrices before and after reordering, respectively) and with fluorescent beads imagery (panels 3,4). Darker pixels depict higher similarity and white pixels depict no similarity. The two red lines in panel 4 indicate a single flip in the computed order that was later corrected with EM imagery. **b**, Volumetric correlative stack of data set 1 with 3 fluorescent channels and 507 consecutive ultrathin sections. Insets: closeups of cell bodies and a neurite carrying different neuroanatomical tracers. The cell bodies in the right panel are outlined with colored dash lines. Blue: tracer injected in Area X. Green: tracer injected in RA. Red: tracer injected in Avalanche. **c**, The EM imagery was connectomics-grade and enabled neurite tracing. Yellow dots: skeletons stemming from 9 seed points placed in a 3×3 grid in the first section of data set 2.

### 5. Imaging

The high packing density of the collected sections on wafer allowed convenient staining procedures (simply exchanging a few microliters of staining solution repetitively on an area smaller than 2 cm x 2 cm), and easy loading into LM and EM microscopes for uninterrupted automated imaging.

After immunostaining against neuroanatomical tracers previously injected into the brain of two zebra finches, multichannel fluorescent imaging (1 and 3 fluorescent channels in datasets 1 and 2, respectively, and 1 widefield channel) was automatically performed with custom scripts. Note that the small wafers were easily coverslipped with mounting medium underneath and oil on top to enable reflection LM with high magnification immersion objectives (63x). After washing off the mounting medium on the wafer followed by heavy metal poststaining, automated scanning EM was performed with custom scripts, acquiring the same portion of the section in each of them (supp. Video 2). Volumetric EM imagery was assembled (contrast enhancement, stitching, affine then elastic alignment) and the LM modality was registered to its EM counterpart (Fig. S7). The whole processing chain was entirely automated with custom scripts^2^ operating in the Fiji/TrakEM2 environment^11,12^. Multibeam scanning EM was also used successfully to image magnetically collected sections (Fig. S5).

### 6. Data analysis

The experiments yielded correlative LM-EM stacks of brain tissue ready for connectomic analysis (Fig. 2b and supp. Video 1). For convenient use, the data were converted to the neuroglancer format and hosted online for seamless browsing and annotation with the web-based tool neuroglancer, also using enhancements for multichannel overlay offered by neurodataviz (Fig. S8). To demonstrate the suitability of the data for connectomic analysis I traced 9 neurites with starting points located on a 3×3 grid within a central area of the first section, Fig. 2c. I also identified structures tagged with an injected neuroanatomical tracer such as an axon making an *en passant* synapse (Fig. S6).

## Conclusion

In conclusion, MagC solves the challenge of collecting hundreds of serial ultrathin sections with a high packing density directly onto silicon wafers. I expect MagC to be used in high-throughput volumetric microscopy beyond connectomics for ultrastructural biology in general. Combined with broad ion beam milling^13^ and next-generation multibeam EM, MagC could become an ideal platform for large-volume EM connectomics.

## Acknowledgements

I thank N. Broguiere and H. Gnaeggi for comments on the manuscript and help in the design of the custom knife boat, respectively. I thank members of the Zeiss MultiSEM team (Eberle, Nickell, Garbowski) for the multibeam scanning EM experiments. I acknowledge support of the Scientific Center for Optical and Electron Microscopy ScopeM of the Swiss Federal Institute of Technology ETHZ.

## Methods

### A. Animal experiments

Animal experiments were approved by the Veterinary office of Canton Zurich (207/2013). Two zebra finches were anesthetized with isoflurane and placed in a stereotaxic device. Fluorescent tracers were bilaterally injected (0.5-1 μL) into different areas^14^ as described in supp. Tables 1,2,3. Three to five days after tracer injection, the animals were sacrificed by perfusion fixation with fixative concentrations of 2% formaldehyde and 2.5% glutaraldehyde in buffer with 0.1M cacodylate, 2mM calcium chloride (referred to as cacodylate buffer). The brain was extracted and slices of 150 µm thickness were cut with a vibratome (Thermoscientific, #Microm HM650V) in cold cacodylate buffer. Portions of the slices containing the nucleus HVC were dissected out with a surgical scalpel and processed similarly as in the protocols described by Deerinck et al.^15^ and Tapia et al.^16^. The sections were washed with cacodylate buffer, stained with heavy metals (2% osmium tetroxide reduced with 1.5% potassium ferrocyanide, washed, 1% thiocarbohydrazide, washed, 2% osmium tetroxide, washed, 1% uranyl acetate at 4C overnight, washed, 0.6% lead aspartate, washed), dehydrated with increasing ethanol concentrations (50%, 70%, 80%, 90%, 95%, 100%, 100%), infiltrated in epoxy Durcupan resin (10g component A/M, 10g B, 0.3g C, 0.2g D), and finally cured in an oven at 52 C for 48 hours.

**Table 1.**
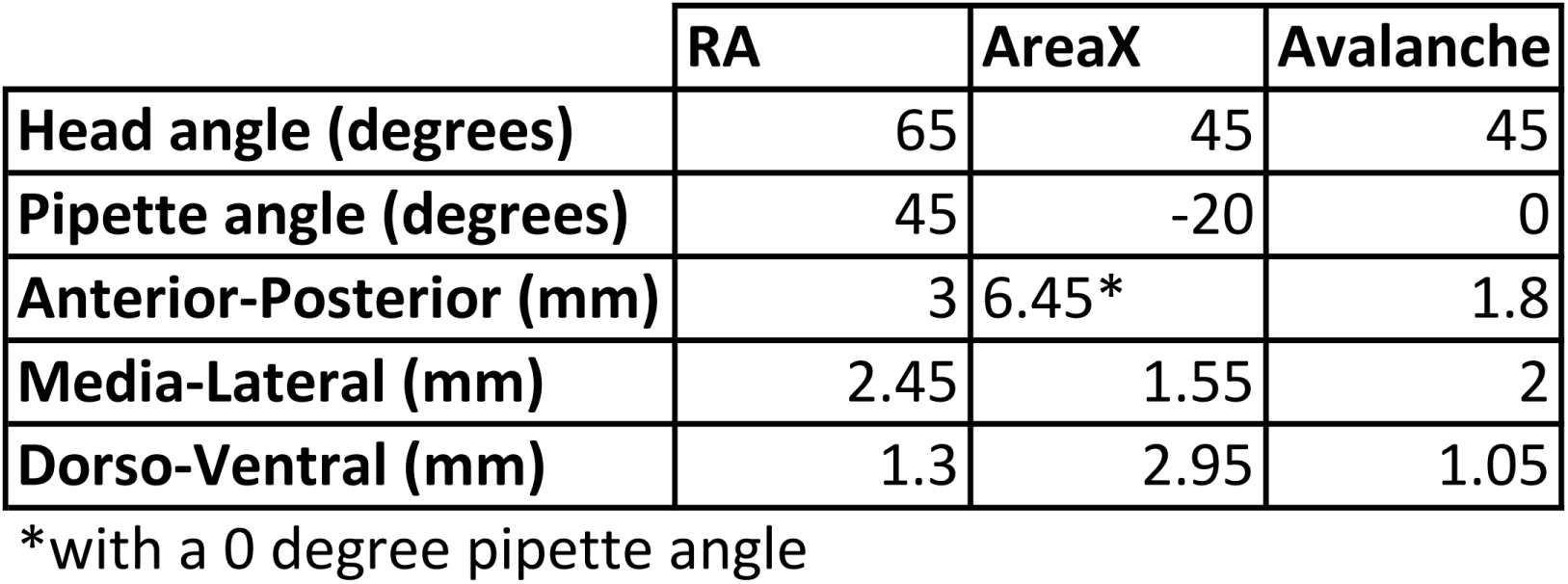
Coordinates of adult male zebra finch nuclei targeted with tracer injections

A. Stereotaxic injection coordinates
|  | RA | AreaX | Avalanche |
| --- | --- | --- | --- |
| **Head angle (degrees)** | 65 | 45 | 45 |
| **Pipette angle (degrees)** | 45 | -20 | 0 |
| **Anterior-Posterior (mm)** | 3 | 6.45* | 1.8 |
| **Media-Lateral (mm)** | 2.45 | 1.55 | 2 |
| **Dorso-Ventral (mm)** | 1.3 | 2.95 | 1.05 |

\*with a 0 degree pipette angle

**Table 2.** Characteristics of the two presented data sets

| Data set | Section number | Anatomical region | Tracer | Injection site | Primary antibody | Secondary antibody | EM size (µm x µm) | EM Dwell time (ns) | Pixel size (nm) |
| --- | --- | --- | --- | --- | --- | --- | --- | --- | --- |
| 1 | 507 | HVC | alexa 488<br>FITC<br>texas red | RA<br>Area X<br>Avalanche | rat α-488<br>mouse α-FITC<br>goat α-rhodamine | 488 α-rat<br>647 α-mouse<br>546 α-rhodamine | 275 x 205 | 820 | 8 |
| 2 | 203 | Dorsal RA | BDA | Caudal RA | mouse α-BDA | 647 α-mouse | 185 x 140 | 6000 | 8 |

**Table 3.** Tracer-antibody library. LT: Life Technologies. VL: Vector Laboratories. JI: Jackson Immunoresearch.

| Antigen | Alexa 488<br>LT #D-22910 | FITC<br>LT #D-1820 | Texas Red<br>LT #D-3328 | BDA<br>LT #D-1956 |
| --- | --- | --- | --- | --- |
| Antibody species | **rabbit** (LT #A-11094)<br>**rat** (Biotem #custom) | **mouse**<br>**rabbit** (LT #A-889) | **goat** (VL #SP-0602) | **mouse** (JI #200-002-211) |

### B. Resin preparation

Magnetic resin was prepared as described by Puig et al.^10^ with 8% weight concentration of iron oxide superparamagnetic nanoparticles (CAN Hamburg, Germany, #SMB-0-038) in epoxy resin (Diglycidylether of Bisphenol A, #D3415 Sigma Aldrich). In addition, fluorescent particles (Cospheric, mean diameter 2 µm, #FMG, #FMR, 0.2% and 1% weight concentration in data sets 1 and 2, respectively) and coumarin dye (SigmaAldrich, #257370, 7-Amino-4-methylcoumarin, 0.5% weight concentration) were added to the resin mixture prior to mixing. The resin mixture was poured between a glass slide (bottom) and a piece of aclar sheet (top), both coated with mould separating agent (#62407445, Glorex, [55]). A PDMS spacer of about 600 μm thickness surrounded the resin and a small weight was put on top of the aclar sheet for flattening. The resin was cured for 6 hours at 70C.

### C. Block augmentation

For block augmentation, a piece of magnetic resin and a dummy were successively glued to the sample of interest using the same Durcupan formulation as described above for brain tissue preparation. The execution details of the procedure are described in Fig. S1.

### D. Section collection

The collection procedure is described in the main text. The custom diamond knife with a bath of dimensions 55 mm x 44 mm (now commercially available, #Ultra ATS, Diatome, Switzerland) was placed in an ultramicrotome (Leica, UC6). The water level in the bath was set with a motorized syringe pump (KDScientific, #210). The setup shown in Fig. 1b consisted of a 3-axis motorized actuator (Thorlabs, #LTS150/M, #PT1/M-Z8) carrying an aluminum plate with a goniometer (Thorlabs, #GN2/M) screwed at its extremity, facing down. A cylindric Neodymium magnet (Supermagnete, cylindrical, 15 mm diameter, 8 mm height) was magnetically anchored to a steel plate screwed to the goniometer. The orientation of the magnet was adjusted with the goniometer in order to make its bottom surface parallel to the water level.

Silicon wafers (Ted Pella, #16015) were cleaved to approximately 40 mm x 45 cm chips, hydrophilized with oxygen plasma (1 min, 25 mA, Emitech #K100X) and placed in the knife bath with a ∼2 degrees angle compared to water level thanks to asymmetrically stacked microscopy coverslips below the wafer chip.

### E. Postembedding stainings

The post-embedding immunostaining protocol is described in the Supplementary methods, paragraph G. Heavy metal post-staining was performed by exposing sections on wafer to a few drops of 2% aqueous uranyl acetate, then to a few drops of Reynold’s lead citrate (lead 4.4% weight concentration), both for 90 seconds. Between the two stains and after the second stain, the entire piece of wafer was immersed consecutively in 3 small petri dishes of double distilled water for 30 seconds each. After the second washing, the wafer was dried with a manual air blower.

### F. Section segmentation

The sections on wafer acquired with low resolution LM (5x air objective, Fig. 1c, S2) were segmented semi-automatically with help of the Trainable Weka Segmentation plugin^17^ in Fiji/TrakEM2 and custom scripts.

### G. Section order retrieval with fluorescent beads

After preprocessing the fluorescent bead imagery (“Normalize local contrast” Fiji plugin, thresholding), the center location of the beads was extracted (Maxima Finder) for each fluorescent channel. The locations of the beads from the two fluorescent channels were merged into a single final channel. I computed a dissimilarity value for every pair of sections. For each pair of bead center sets, descriptor matching was performed (using descriptor-based bead alignment available in Fiji^18^). If no geometric match was found for a given pair of sections, then the dissimilarity value was set to a fixed large number. If a geometric match was found, then a matching affine transform was computed and applied to the first bead set, thus bringing the pair of bead sets into a same coordinate system. In this common coordinate system, the bead centers contained outside a central bounding box were excluded from further calculations to avoid considering beads that are present in one section but not in the other one due to a limited field of view and due to the different orientations of the section. The pair of remaining bead sets was then matched again with the descriptor-based tool. For each match, that is each pair of two matching beads, the absolute difference of the diameters of the matching beads was computed. The dissimilarity of two sections was then defined as the sum of these diameter differences across all matching beads. A traveling salesman problem was formulated using the dissimilarities as distances between nodes of a graph, and the problem was solved with the Concorde solver^19^.

### H. Section order retrieval with EM

An EM section was made of a mosaic of EM tiles (3×3 or 2×2). For a given pair of EM sections, a dissimilarity was computed for each pair of corresponding mosaic tiles and averaged across the tiles to yield the complete dissimilarity between two EM sections. The dissimilarity of two tiles was calculated as follows: an affine transform matching was sought between the pair of images, using the SIFT matching algorithms implemented in Fiji. If no affine transform was found, then the pair of tiles was given an arbitrary high dissimilarity. If a transform was found, then it was used to align the two tiles and a normalized cross-correlation was computed in a central box of 2000 x 2000 pixels. The value (2 – correlation) was used as the dissimilarity value between the two tiles. When averaging the dissimilarities across tiles for a given pair of EM sections, the non-matching tiles were excluded if other tiles were matching. It made the dissimilarity value more robust to artefacts that may have prevented a match to be found in one of the tiles. As with the beads, an open traveling salesman problem was solved with the computed dissimilarities and yielded the original order, as confirmed with manual inspection of the EM stack.

### I. Imaging

Widefield fluorescent LM and scanning EM were performed with the characteristics detailed in Supp. Table 2. The LM and EM were controlled with python scripts through micromanager^20^ and the Zeiss API, respectively. Autofocus was performed on each tile in LM (Nikon, #PerfectFocusSystem) and at the center of each EM mosaic.

### J. Data assembly

The brightfield channel of the LM imagery was used for the stitching, alignment and registration operations (with an initial “Normalize local contrast” from Fiji with blocks of about 100 pixel x 100 pixel) done in Fiji. The stitching was then propagated to all fluorescent channels. Stitching and alignment^21^ of the EM imagery was done with custom scripts in TrakEM2.

For cross-modality registration, the stitched mosaics of the brightfield channel were preprocessed with local contrast enhancement and Gaussian blurring. The EM counterpart mosaics were downsampled to exhibit roughly the same pixel size as the LM imagery and further preprocessed with local contrast enhancement. The LM brightfield and EM imageries then exhibited a similar appearance so that corresponding SIFT features^22^ could be computed across the two modalities (Fig. S7). Moving least squares transforms^23^ were computed based on these matching SIFT features using Fiji. The transforms were then upsampled and applied to all fluorescent channels of the LM imagery in the TrakEM2 plugin to yield a volumetric correlative LM-EM stack.

For visualization purposes, the correlative LM-EM imagery was converted to the neuroglancer format and hosted online for convenient in-browser visualization and annotation. Details of the procedure in supp. Text 1 (paragraph N.). I plan to make data available in *neurodata.io*. The data sets 1 and 2 are currently available for review at. and ., respectively.

## Supplementary

### A. Stereotaxic injection coordinates

### B. Characteristics of the two datasets

The color coding of the fluorescent imagery in data set 1 is: Green - tracer injected in RA, Red-tracer injected in Avalanche, Blue-tracer injected in Area X.

### C. Block augmentation

**Figure S1.**
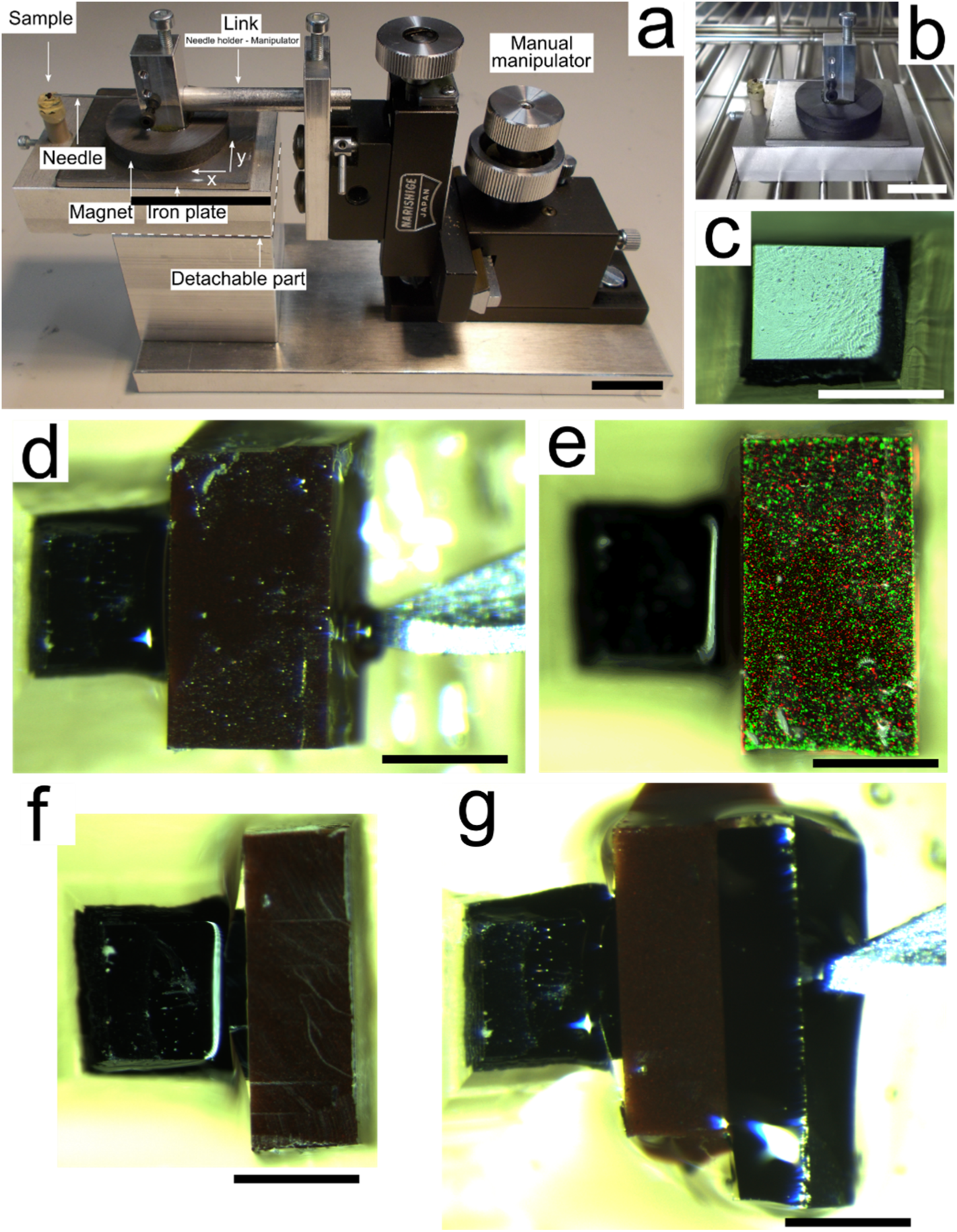
Magnetic augmentation. **a,** Mounting helper device. The manual manipulator allows the experimenter to precisely place the needle in contact with the block to be glued on the sample. The iron plate together with the base magnet of the needle holder maintain the position set by the manipulator. **b,** The detachable part is placed in an oven for temperature curing. **c**, The biological sample is trimmed manually with a razor blade. **d,** The magnetic resin is glued to the sample and maintained in place with a needle. **e,** Overlay of color and fluorescent imagery showing the fluorescent particles contained in the magnetic resin. **f**, The magnetic resin is manually trimmed down to achieve roughly a 50/50 ratio of sample surface to magnetic surface suitable for sectioning at 50 nm nominal thickness. **g,** An additional dummy piece of heavy metal-stained resin embedded brain tissue is glued to the block.h After final trimming, the block presents a pointed shape (overlayed with the yellow dashed line) and is ready for sectioning. Note: the stronger red signal at the bottom is not due to large particle aggregates but due to out-of-focus effects. Scale bars: **a**,**b**-20 mm; **c** to **g**: 500 μm

### D. Wafer and section overview

**Figure S2.**
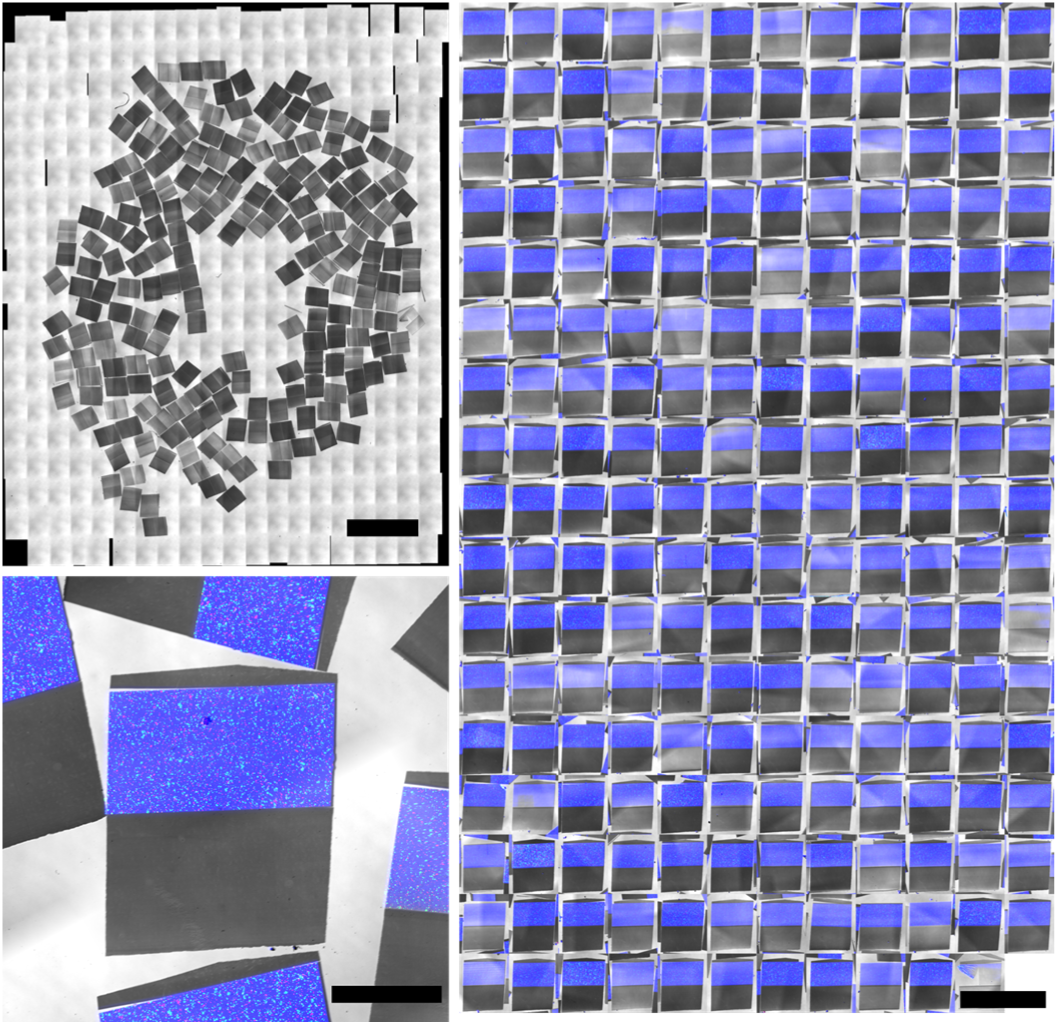
Wafer overview of data set 2. **Top left**, widefield LM of the collected sections. **Bottom left**, close up around one section, merge multichannel fluorescent imagery: blue-Coumarin, green,red-fluorescent beads). **Right**, montage of all sections with the same orientation.

### E. Metric for order retrieval and order retrieval for data set 1

A metric was defined to assess the quality of the reordering process based on imagery of fluorescent beads. This metric requires the knowledge of the ground truth order, which I obtained from the SOR performed with EM imagery, and which I call the “EM order”.

For each section of a reordered dataset, a cost is given to the the link between the given section and the next one. The cost is equal to the difference of the indices of the sections in the ground truth order, minus one. For example, the links of the order 1-2-3-4-5-6-7-8 have the costs 0,0,0,0,0,0,0, so do the links of the 8-7-6-5-4-3-2-1 order, while the links of the order 1-2-4-5-3-8-6-7 have the costs 0,1,0,1,4,1,0. A single flip such as 1-**2**-**4**-3-5-6 has the cost 0,1,0,1,0,0. The frequency of these costs gives an estimate of how precise the reordering is.

**Figure S3.**
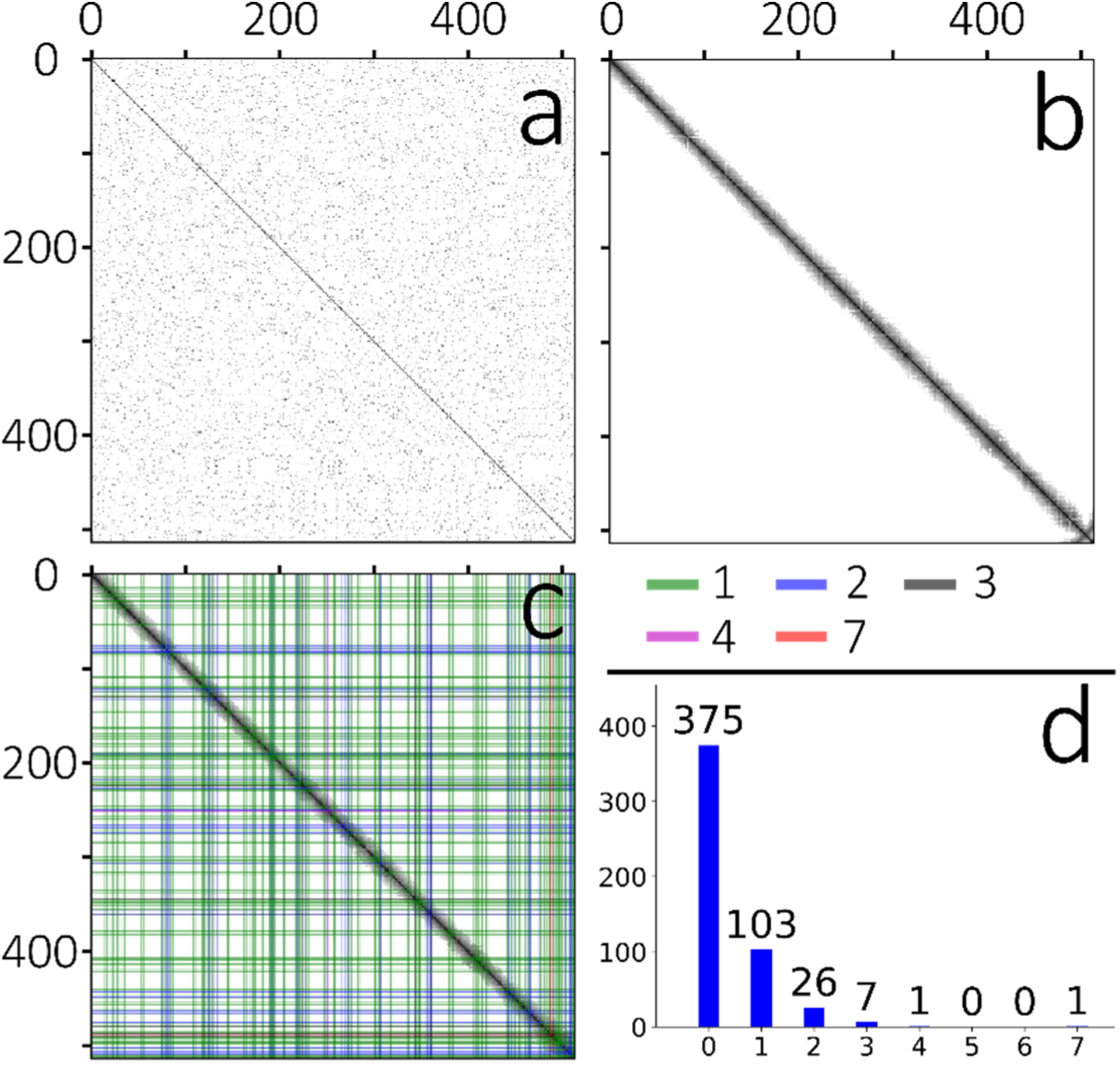
Section order retrieval of data set 1 (low concentration of beads). **a,** Matrix of pairwise similarities of unordered sections computed with EM imagery. Darker pixels depict higher similarity and white pixels depict no similarity. **b,** Reordered EM matrix. **c**, Matrix of pairwise similarities of unordered sections computed with fluorescent bead imagery. The original order is the order provided by the section segmentation pipeline. **d**, Similarity matrix of the reordered sections. The order overall looks consistent, except slight deviations at the end of the data set (around section number 500) that can be seen in the lower left of the matrix. **e**, Similarity matrix with the EM order. The costs of the links of the bead order are overlayed as vertical and horizontal bars of different colors. The green bars show the locations of the links that have a cost of 1, and there are 103 of them. **f,** The distribution of costs of the links. The largest mistake is a link with cost 7 at the end of the data set.

### F. EM of magnetic resin

**Figure S4.**
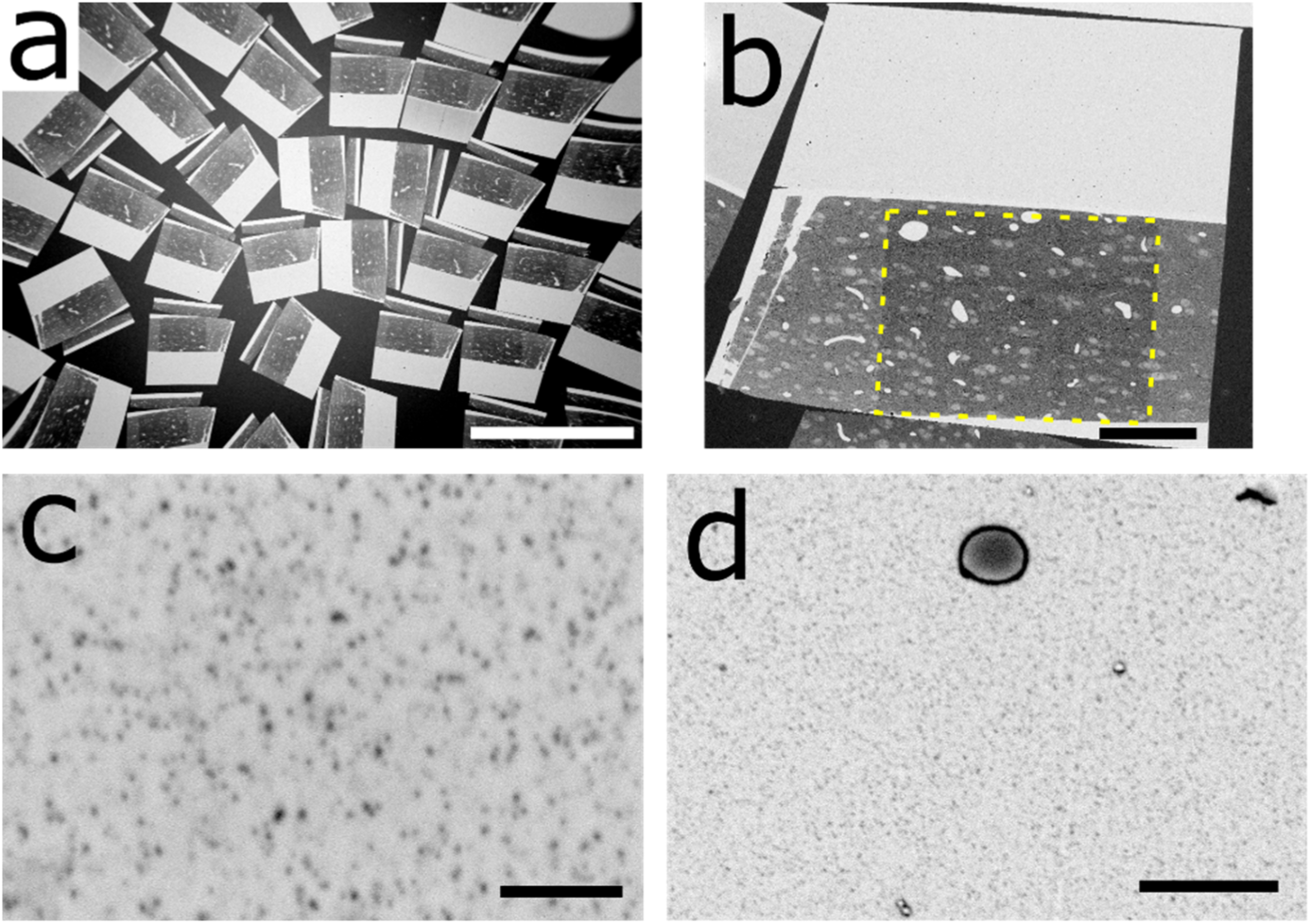
Electron micrographs of the sections from data set 1. **a,** Electron micrograph of numerous sections collected on wafer. **b**, EM of a section. The yellow dashed square highlights the region that has been imaged with the electron microscope and became darker due to the beam irradiation. **c**, EM of well-dispersed superparamagnetic nanoparticles in the appended resin. **d**, Small contaminations can sometimes be found in the fluomagnetic resin. Scale bars: a-1mm, b-100 µm, c-500 nm, d-2µm

### G. Immunostaining protocol

I deposited and exchanged staining solutions manually with graduated pipettes on the sections collected on flat substrate. All steps were performed at room temperature. The blocking solution was: 1% Baurion BSA-c, 0.05% Tween^24^ in TBS pH 7.4. The detailed procedure was:

1. Blocking -- blocking solution -- 2x 10 min
2. Primary antibody incubation -- 1:50 in blocking solution -- 1.5h
3. Washing -- TBS -- 4×5min
4. Secondary antibody -- 1:100 in blocking solution -- 1 h
5. Washing -- TBS -- 2×5min
6. Washing -- dH2O -- 2×5min
7. Drying with hand dust blower (Bergeon #30540) 8. Air drying – 5 min

Proceed to fluorescent imaging within the next hours to avoid decay of staining as reported by Micheva et al. ^25^, Fig. Sup. S3.

### H. Multibeam 294 scanning EM

**Figure S5.**
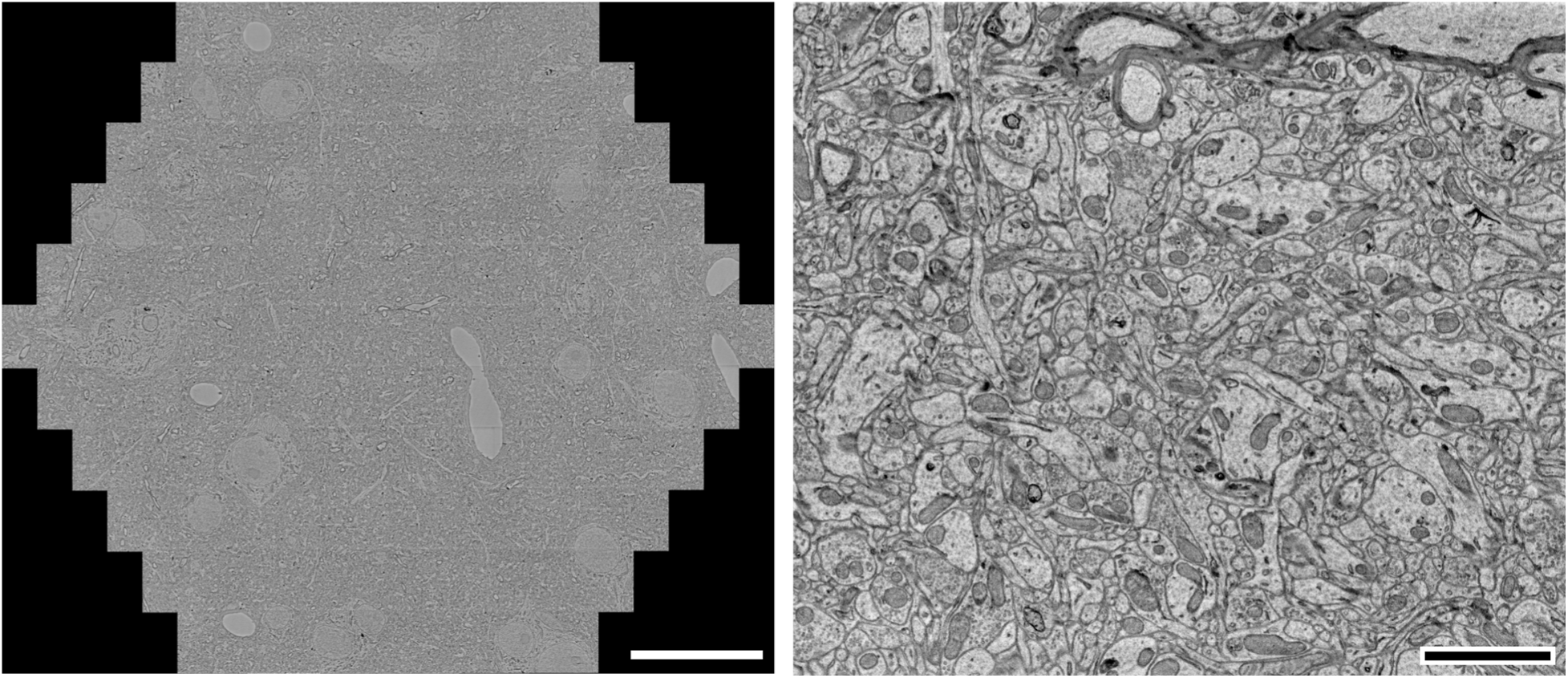
Multibeam (91) Scanning EM of magnetically collected sections on silicon wafer. **Left**, Overview of 91 stitched tiles. **Right**, a tile produced by one of the 91 beams. Imaging conditions: silicon wafer chip glued to EM stub with carbon glue, 4 nm pixel size, 400 ns dwell time, Scale bars: **left**, 20 μm; **right**: 2 μm

### I. Labeled neurite

**Figure S6.**
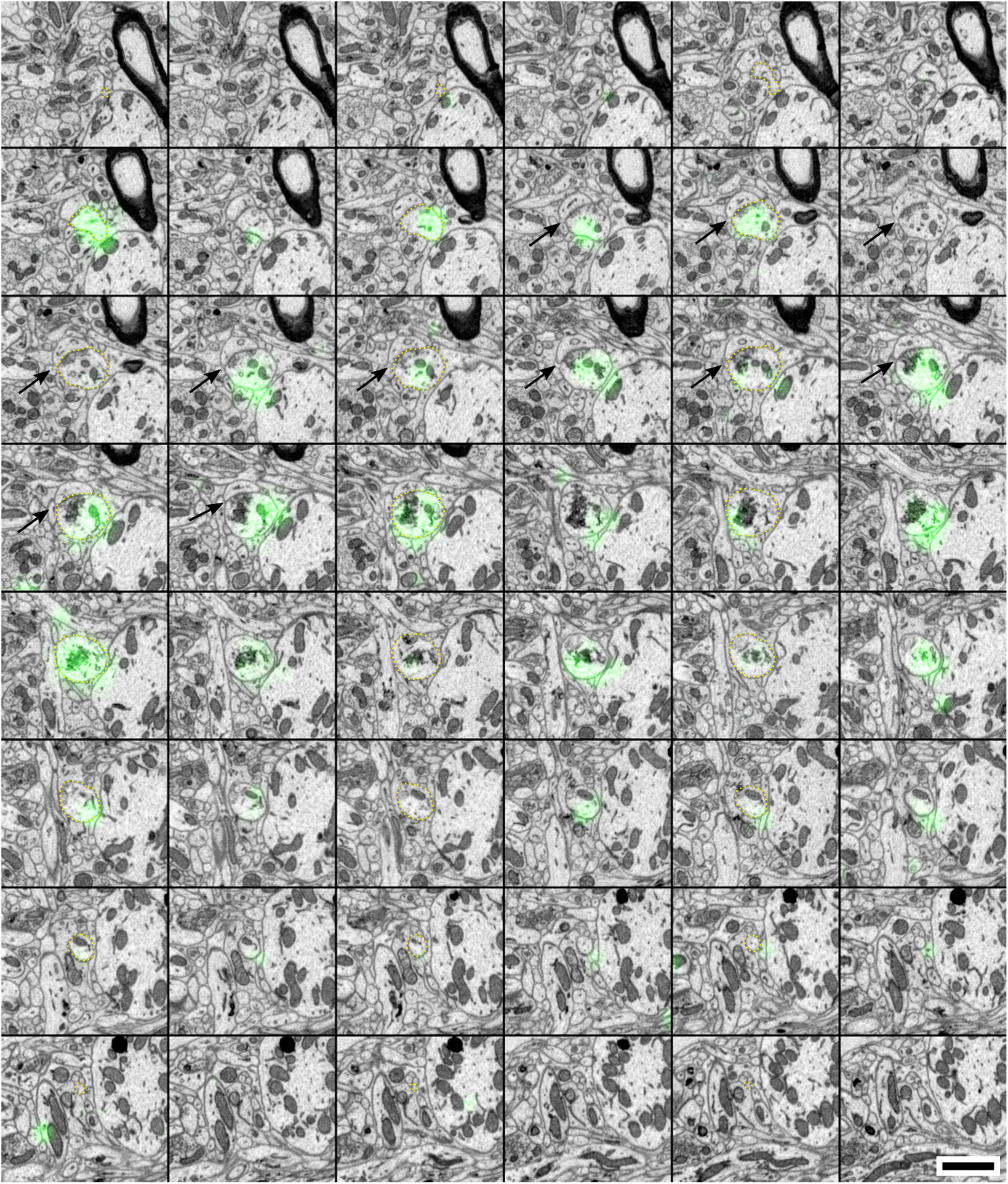
Labeled axon makes a synapse en passant. The axon is delineated with dashed yellow lines (every second section). Black arrows indicate the synapse. Scale bar 1 micron.

### J. Cross-modality registration

**Figure S7.**
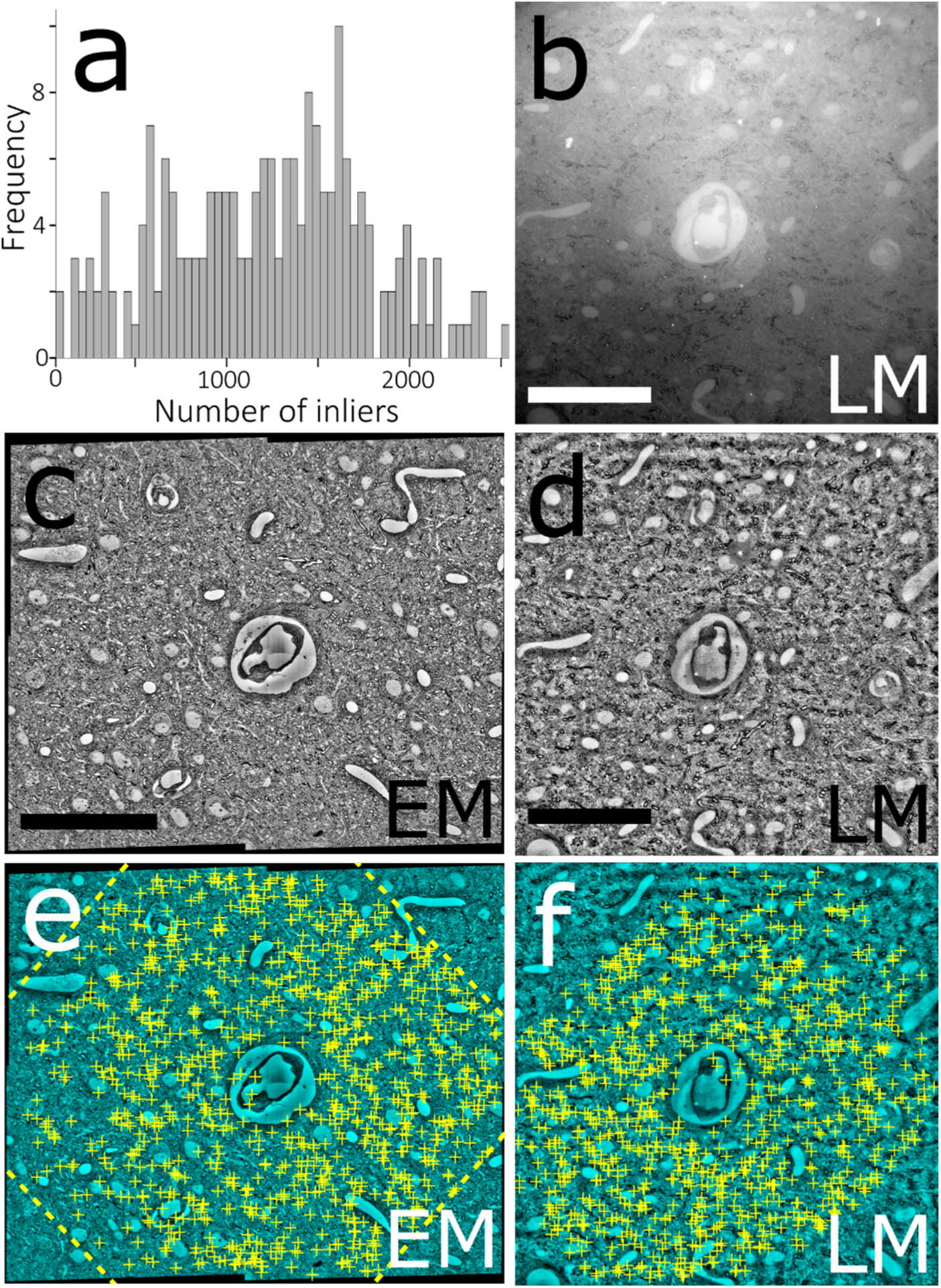
Automated LM-EM registration. **a**, Histogram of number of matching inliers found for each of the 203 LM-EM pairs of data set 2. **b**, A reflection brightfield light micrograph after simple thresholding. **c**, downscaled EM mosaic. **d**, Same micrograph as in a after local contrast normalization. Note the high similarity with its EM counterpart micrograph in c. **e,f**: Same micrographs as in c,d, respectively. The yellow crosses show the location of matching SIFT features between the two images. The dashed yellow lines in e show the outline of LM micrograph when affine transformed to match its EM counterpart. Scale bars: 50 μm

### K. Video 1: flythrough in EM imagery of data set 2

Video available here: https://youtu.be/VL0F9DkZVaQ

### L. Video 2: zoom on wafer of data set 2

Video available here: https://youtu.be/23DWA8YbGH4

### M. Visualization of correlative LM-EM stack in neuroglancer

**Figure S8.**
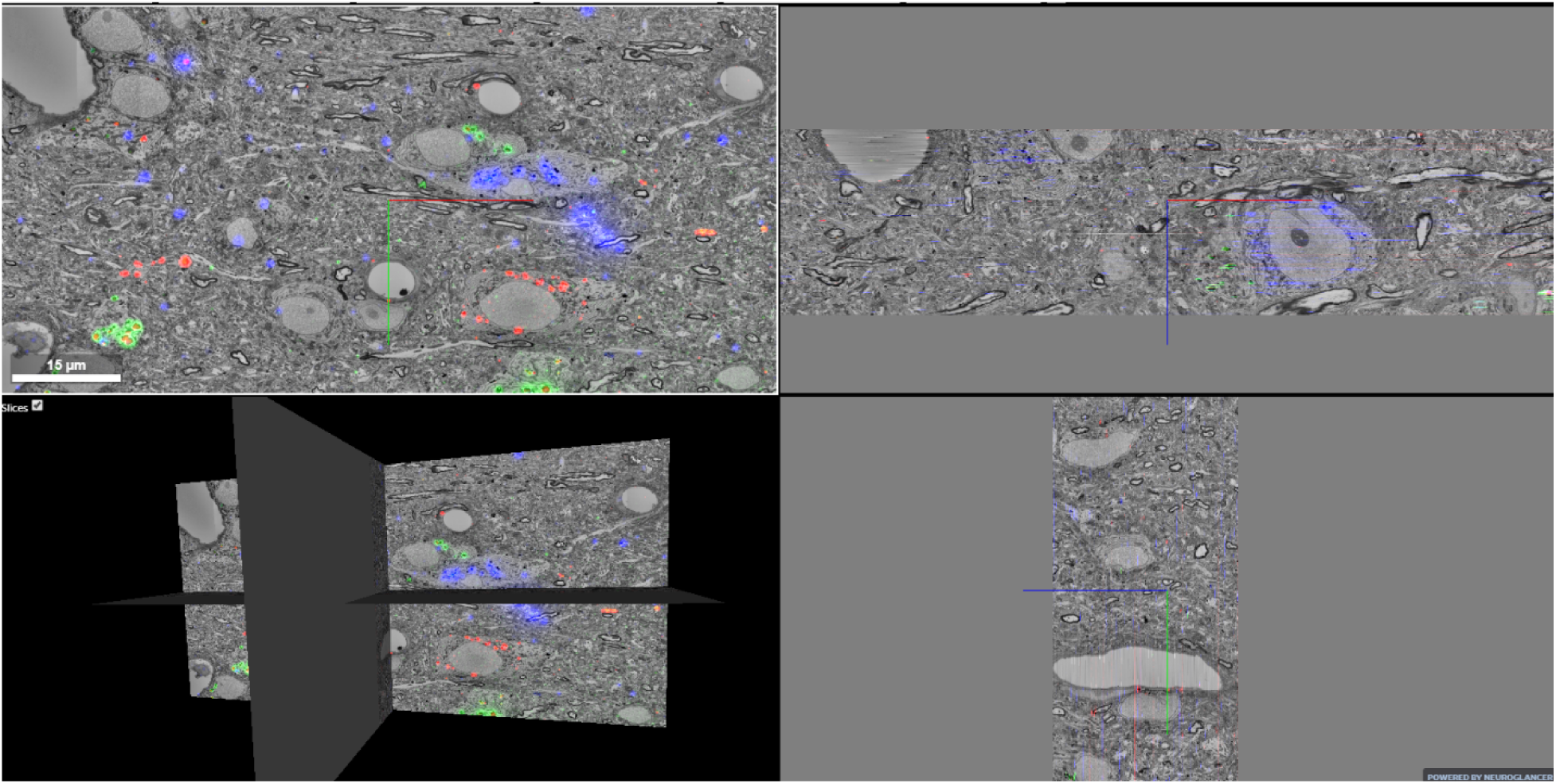
Multicolor correlative LM-EM imagery (data set 1) of zebra finch HVC nucleus showing 3 neuroanatomical tracers injected in Area X (blue), the nucleus Robustus of the Arcopallium (green), and Avalanche (red). The two panes on the right show x and y reslices through the volume.

### N. Text 1: conversion of correlative LM-EM imagery for neuroglancer

EM imagery assembled in TrakEM2 along with all transforms (affine, elastic and moving least squares) was converted into a Render^3^ project^26^ with custom scripts and the TrakEM2 converter script of the Render project. Similarly, TrakEM2 projects were created for each LM channel that contained stitching and moving least square transforms. These TrakEM2 projects were converted to separate Render projects. The imagery of the EM and LM Render projects was rendered to files using a custom script and the Render script for mipmap creation (render_catmaid_boxes). With a custom script, these mipmaps were then used to create chunks at different resolutions in the “precomputed format” of Neuroglancer^4^. The chunks were uploaded to an online cloud storage service (Google storage) and an instance of the Neuroglancer software hosted online (neurodataviz from the MICrONS project) was used to visualize the data. The EM imagery and each fluorescent LM channel were added into a neuroglancer session as separate data sources. After online visualization with neuroglancer, stacks of correlative imagery were fetched using the cloud-volume library^5^. Neurite tracings were performed in neuroglancer (line annotations).

### O. Traced neurites

The 9 traced neurites in data set 2 are available online at this link (copy-paste in browser address):

## Footnotes

2 https://github.com/templiert/MagC

3 https://github.com/saalfeldlab/render

4 https://github.com/google/neuroglancer

5 https://github.com/seung-lab/cloud-volume

